# Natural selection promotes the evolution of recombination 1: between the *products* of natural selection*

**DOI:** 10.1101/2021.06.07.447320

**Authors:** Philip J Gerrish, Benjamin Galeota-Sprung, Paul Sniegowski, Alexandre Colato, Julien Chevallier, Bernard Ycart

## Abstract

Shuffling one’s genetic material with another individual seems a risky endeavor more likely to decrease than to increase offspring fitness. This intuitive argument is commonly employed to explain why the ubiquity of sex and recombination in nature is enigmatic. It is predicated on the notion that natural selection assembles selectively well-matched combinations of genes that recombination would break up resulting in low-fitness offspring – a notion often stated in the literature as a self-evident premise. We show however that, upon closer examination, this premise is flawed: we find to the contrary that natural selection in fact has an encompassing tendency to assemble selectively mismatched gene combinations; recombination breaks up these selectively mismatched combinations (on average), assembles selectively matched combinations, and should thus be favored. The new perspective our findings offer suggests that sex and recombination are not so enigmatic but are instead unavoidable byproducts of natural selection.

## I. INTRODUCTION

It seems recombination should be disadvantageous most of the time. High-fitness genotypes that are amplified by natural selection (*products* of natural selection), it seems, should carry “good” (selectively well-matched) combinations of genes. And recombination, which shuffles genes across individuals, should only break up these good combinations and thus should be evolutionarily suppressed. In this light, the overwhelming prevalence of recombination across the tree of life is a mystery.

The foregoing paragraph outlines a line of reasoning commonly employed to demonstrate why the ubiquity of sex and recombination is enigmatic. The premise of this line of reasoning – that natural selection will tend to amplify genotypes carrying “good” (selectively well-matched) combinations of genes – is so intuitive that it is considered self-evident in much of the literature [3–12] and has largely gone unquestioned.

We define a *product* of natural selection to mean any genotype that has become locally prevalent at any scale – e.g., population, subpopulation, deme, niche, competing clone, etc. – through the local action of natural selection.

Recombination can only have an effect on offspring fitness if the genetic makeup of the parents differ. In a structured population, local evolution can lead to divergence in genetic makeup. If two parents come from two different locally-evolved subpopulations, therefore, they will likely differ in their genetic makeup. The question then becomes, will they differ in such a way that tends to enhance or diminish the fitness of their offspring? In other words, will the offspring of different *products* of natural selection (as defined above) tend to be superior or inferior to their parents? The standard argument outlined in the first paragraph would imply that offspring fitness should be inferior to parent fitness, because recombination would break up good gene combinations that each parent had acquired in their local environments.

An answer to this question came early on from agriculture. That the out-crossing of inbred lineages tends to confer *vigorous* offspring (later dubbed “hybrid vigor” or “heterosis” [13–16]) is an observation that has likely been part of farmer folklore for centuries. The earliest known systematic study of this phenomenon was conducted by Darwin himself [17, 18]; his study was perhaps motivated, at least in part, by his search for a theory of inheritance that was consistent with his theory of natural selection [18, 19]. Observations of heterosis gave his “blending” theory of inheritance a plausible foothold: either chance differences in the founding individuals of two locally-evolving subpopulations or divergent selection pressures in these subpopulations would give rise to persistent genetic differences across subpopulations despite local blending within each subpopulation.

Translated to the language used in the present study, what Darwin was documenting in these early studies was that recombination between two *products* of selection tends to produce high-fitness offspring (assuming that fitness and “vigor” are correlated). This observation contradicted Darwin’s theory of blending inheritance which predicts offspring fitness at the midpoint between parent fitnesses; heterosis, it would seem, had the potential at least to reveal the flaw in his theory.

That these early studies of inbreeding and heterosis are relevant to the evolution of sex and recombination is not a new idea [11, 20, 21], and is subsumed under Lewontin’s general proclamation that “every discovery in classical and population genetics has depended on some sort of inbreeding experiment” [20, 22, 23]. Since these early studies, several more recent studies have shown that different kinds of population structure can create conditions that make recombination across locally-evolving subpopulations favorable [10–12, 24–29]. These studies find that population structure can help to maintain the variation without which recombination would be neither advantageous nor disadvantageous, and they identify conditions under which recombination is advantageous. Spatially heterogeneous selection can, for example, create negative fitness associations if selection acts more strongly on one gene (one *locus*) in one spatial “patch” and acts more strongly on a different locus in a neighboring patch [24, 25, 29]. The negative fitness associations that arise in such a scenario would be broken up by recombination thus giving recombination-competent (*rec*^+^) lineages a selective advantage. The prevalence of such scenarios in nature, however, is unknown and questionable [25].

Generally speaking, recombinants whose parents are two distinct products of natural selection will carry an immediate selective advantage, on average, when the ensemble of such products harbors an excess of selectively antagonistic (mismatched) gene combinations and a deficit of synergistic (well-matched) combinations: by randomly shuffling different gene variants (or *alleles*) across different products of selection, recombination will on average increase offspring fitness. The challenge in explaining the ubiquity of sex and recombination in nature is to identify a source of this selective imbalance that is comparably ubiquitous. One feature of living things whose prevalence approximates that of sex and recombination is evolution by natural selection. In the present study, we assess the effects of natural selection by itself on selective imbalance among products of selection. In doing so, we determine the selective value of recombination in structured (e.g., spatially structured) populations. We find that natural selection by itself has an encompassing tendency to amplify selectively mismatched combinations of alleles, thereby promoting the evolution of recombination across different products of selection.

## II. MEASURING SELECTIVE IMBALANCE

In much of the relevant literature, the measure of selective mismatch across loci affecting the evolution of recombination is *linkage disequilibrium* (LD) [26, 30–35], which measures the covariance in allelic *states* across two loci [36] (i.e., it measure the bias in allelic frequencies across loci) but does not retain information about the selective value of those alleles.

For the sake of presentation, we here consider an organism with just two fitness-related genes (or two *loci*) whose fitness contributions are represented by random variables *X* and *Y*. We have found (see ev2 [2]) that the expected selective advantage of newly-formed recombinants (and the advantage of recombination over the course of a single generation) is

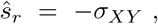

where *σ*_*XY*_ is the covariance between *X* and *Y*. This measure of selective imbalance is superior to LD in that it retains information about both the frequencies and selective value of alleles and it directly gives the selective advantage of recombinants. Furthermore, we show in ev2 [2] that *ŝ*_*r*_ defined in this way provides a lower bound for the selective advantage of a *rec*^+^ lineage within a single population. Our results will thus be given in terms of covariance.

## III. NATURAL SELECTION: SIMULATIONS

As an introduction to how we model the selective value of recombination across different products of selection, we begin by describing simple simulations. We encourage interested readers to perform these very simple simulations to see for themselves the counter-intuitive outcome and its remarkable robustness to the choice of distribution.

In order to isolate the effects of natural selection, we assume the population size to be infinite so that dynamics are deterministic (as stated in companion studies [1, 2]). As we will show later, however, our findings are fairly robust to relaxation of this assumption. We will assume the organism in question has two loci. The simulations begin by generating a set of *n* distinct genotypes; this is achieved simply by drawing *n* genic fitness pairs (*x*_*i*_, *y*_*i*_), *i* = 1, 2, …, *n* at random from some bivariate distribution. The bivariate distribution can be any distribution with any covariance.

Next, the simulation simply records the (*x*_*i*_, *y*_*i*_) pair whose sum *x*_*i*_ + *y*_*i*_ is the largest and puts this pair into a new array that we will denote by 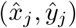. This mimics natural selection acting in an infinite population; in an infinite population there is no role for chance and natural selection thus deterministically fixes the fittest genotype.

The procedure is then repeated a few thousand times, so that there are a few thousand entries in the 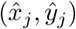 array of “winners”, or “products of selection”. The covariance of the 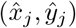 array is then computed. Remarkably, this covariance is always less than the covariance of the initial bivariate distribution used to generate the (*x*_*i*_, *y*_*i*_). If the covariance of the initial bivariate distribution is zero (i.e., if *X* and *Y* are independent), the covariance between *X* and *Y* among the “products of selection” will always be negative (i.e., the mean value of recombinants across different products of selection will always be positive). The interested reader may want to explore this case first, because: 1) she/he will see that any bivariate distribution from uniform to Cauchy gives this result, and 2) this is the case that is the primary focus of the following mathematical developments. An example set of such simulations where *X* and *Y* are skew-normal is plotted in Fig 1.

**Figure 1.**
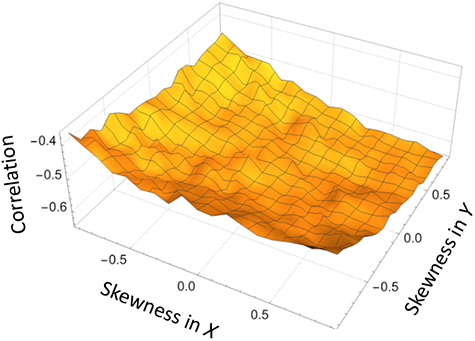
Correlation between genic fitness 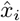 and *ŷ*_*i*_ among products of selection in simple simulations. A set of 20 *x*-values was drawn from a skew-normal distribution with mean −0.1, standard deviation 0.1 and skewness indicated by the *x*-axis. A set of 20 *y*-values was drawn from a skew-normal distribution with mean −0.1, standard deviation 0.1 and skewness indicated by the *y*-axis. These *x* and *y* values were paired up to form an array of 20 (*x, y*) pairs. The pair whose sum *x* + *y* was the largest was selected and its values appended to a new array 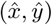 of products of selection. This was repeated 5000 times. The correlation between 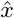 and *ŷ* was computed and plotted for each pair of skewness values.

## IV. NATURAL SELECTION: ANALYSIS

We now turn to mathematical analyses of the procedure described above for simulations. We begin with a generalization of what we describe above: here, instead of two loci with genic fitnesses *x*_*i*_ and *y*_*i*_ for the *i*^*th*^ genotype, we have *m* loci and a vector of genic fitnesses (*x*_*i*1_, *x*_*i*2_, …, *x*_*im*_). Next, we zero in on analyses of the simplest scenario of two loci and two genotypes. Extrapolation of our qualitative results from this simplest-case scenario to more loci and more genotypes is corroborated by simulation (SM). As will become apparent, the mathematical analyses eventually require some restrictions on the bivariate distribution governing genic fitnesses in the initial population. Simulations, however, show that our findings hold qualitatively for essentially any distribution chosen.

### General setting: *m* loci, *n* alleles

Let *n* and *m* be two positive integers. Let (*X*_*i,j*_)_1⩽*i*⩽*n*;1⩽*j*⩽*m*_ be a rectangular array of independent random variables. For our purposes, each *X* quantifies a fitness-related phenotype encoded at one locus. Each row represents an individual’s haploid genome and each column represents a locus on that genome. See Fig. 2. We shall denote by *X*_*i*_ = (*X*_*i,j*_)_1⩽*j*⩽*m*_ the *i*-th row of the array (the *i*-th individual in a population). Let *ϕ* be a measurable function from ℝ^*m*^ into ℝ. For *i* = 1, …, *n*, denote by *Z*_*i*_ the image by *ϕ* of the *i*-th row of the array.

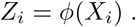

**Figure 2.**
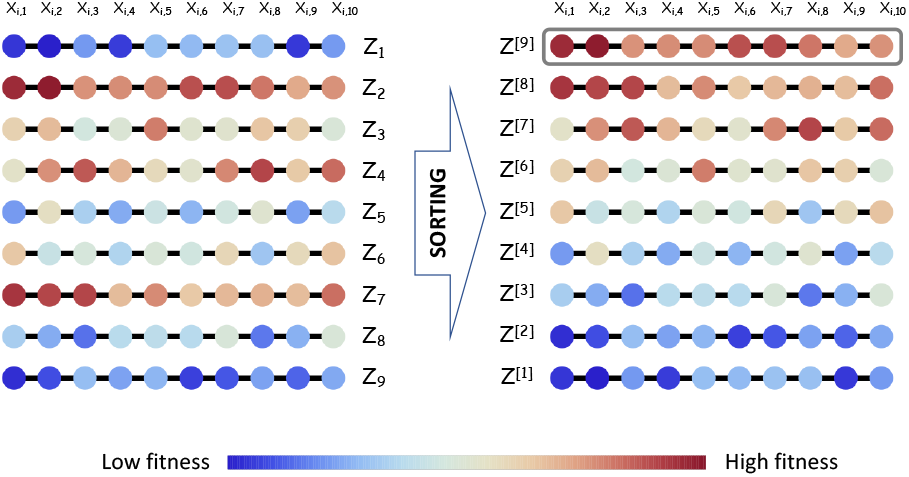
General setting. The population here consists of *n* = 9 genotypes represented by the 9 rows, each of which carries a genome with *m* = 10 loci represented by the 10 columns. Each dot represents a locus on an individual genome and its color indicates its genic fitness. The total fitness of the *i*^*th*^ individual is *Z*_*i*_ = *ϕ*(*X*_*i*1_, *X*_*i*2_, …, *X*_*im*_), where *X*_*ij*_ is the genic fitness of *j*^*th*^ locus in the *i*^*th*^ genotype. Strictly speaking, *ϕ* can be any increasing function of the genic fitnesses, *X*_*ij*_. To give a simple and useful example, *ϕ* may be defined simply as the sum of its arguments. We employ this definition of *ϕ* extensively in the main text and in our analyses, both because of its simplicity and because of its connection to classical population genetics and notions of additive fitness. On the left-hand side, the genomes are not sorted in any order; on the right-hand side, the same genomes are sorted (ranked) by their total fitness, *Z*, such that *Z*^[1]^ is the genome of lowest fitness and *Z*^[*n*]^ is the genome of highest fitness. In an infinite population (deterministic selection), the fittest genome (*Z*^[*n*]^, highlighted by a frame) always eventually displace all other genomes. The statistical properties of the genic fitnesses of this fittest genome are thus of special interest from an evolutionary perspective. In particular, we are interested in any statistical associations among these genic fitnesses: if that association tends to be negative, then recombination will be favored.

*Z*_*i*_ represents the total fitness of genotype *i*. Denote by *σ* ∈ *𝒮*_*n*_ the random permutation such that

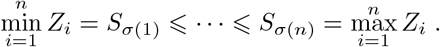

The permutation *σ* is uniquely defined up to the usual convention of increasing order for indices corresponding to ties. Deterministically, natural selection will cause the genome of highest fitness 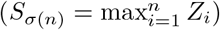 to fix. We are interested in the statistical properties of the *X*_*σ*(*n*),*j*_ ; in particular, we are interested in any associations that might arise across loci (across different values of *j*) in this winning genotype. If these associations are negative, recombination – which alleviates negative associations across loci – should be favored.

For 1 ⩽ *i* ⩽ *n* and 1 ⩽ *j* ⩽ *m*, define:

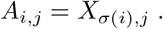

For 1 ⩽ *i* ⩽ *n, A*_*i*_ = (*A*_*i,j*_)_1⩽*j*⩽*m*_ is that row in the array (*X*_*i,j*_) which ranks *i*-th in the order of images by *ϕ*.

#### Density

##### Proposition 0.

*Assume that for j* = 1, …, *m, X*_*i,j*_ *has pdf f*_*j*_, *for all i* = 1, …, *n. Denote by H the common cdf of the Z*_*i*_*’s and assume that H is continuous over its support. The joint pdf of A*_*i*_ *is:*

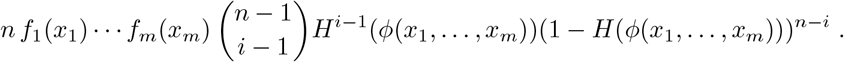

*Proof*: For any continous bounded function Ψ of *m* variables:

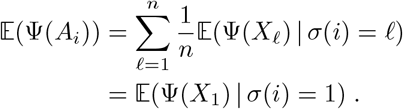

Thus the distribution of *A*_*i*_ and the conditional distribution of *X*_1_ given that Φ(*X*_1_) ranks *i*-th, are the same. The pdf of *X*_1_ is *f*_1_(*x*_1_) … *f*_*m*_(*x*_*m*_). The probability of the event *σ*(*i*) = 1 is 1*/n*. Conditioning on *X*_1_ = (*x*_1_, …, *x*_*m*_), the probability that *X*_1_ ranks *i*-th is the probability that among *Z*_2_, …, *Z*_*n*_, *i* − 1 are below *ϕ*(*x*_1_, …, *x*_*m*_) and *n − i* are above. The probability for *S*_𝓁_ to be below *ϕ*(*x*_1_, …, *x*_*m*_) is *H*(*ϕ*(*x*_1_, …, *x*_*m*_)). Hence the result. □

Observe that the average of the densities of *A*_*i*_ is the common density of all the *X*_*i*_, *i*.*e. f*_1_(*x*_1_), …, *f*_*m*_(*x*_*m*_). This was to be expected, since choosing at random one of the *A*_*i*_ is equivalent to choosing at random one of the *X*_*i*_. The question is whether the *A*_*i*_ are negatively associated in the sense of Joag-Dev and Proschan [37]; this seems a reasonable conjecture in light of Theorems 2.8 and also examples (b) and (c) of section 3.2 in that reference.

### Two loci, two alleles

No hypothesis on the ranking function *ϕ* is made at this point, apart from being measurable. Notations will be simplified as follows: (*X*_1_, *Y*_1_, *X*_2_, *Y*_2_) are i.i.d.; (*X*_(1)_, *Y*_(1)_) (the *infimum*) denotes that couple (*X*_1_, *Y*_1_) or (*X*_2_, *Y*_2_) whose value by *ϕ* is minimal; (*X*_(2)_, *Y*_(2)_) (the *supremum*) denotes that couple (*X*_1_, *Y*_1_) or (*X*_2_, *Y*_2_) whose value by *ϕ* is maximal.

#### Proposition 1.

*Let ψ be any measurable function from* ℝ^2^ *into* ℝ. *Then:* 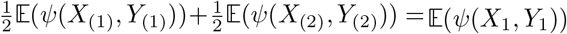. *In particular, the arithmetic mean of* 𝔼(*X*_(1)_) *and* 𝔼(*X*_(2)_) *is* 𝔼(*X*_1_).

*Proof*: Consider a random index *I*, equal to “(1)” or “(2)” each with probability 1*/*2, independent from (*X*_1_, *Y*_1_, *X*_2_, *Y*_2_). By an argument used in the previous section, the couple (*X*_*I*_, *Y*_*I*_) is distributed as (*X*_1_, *Y*_1_). Hence, 𝔼(*ψ*(*X*_*I*_, *Y*_*I*_)) = 𝔼(*ψ*(*X*_1_, *Y*_1_)), however,

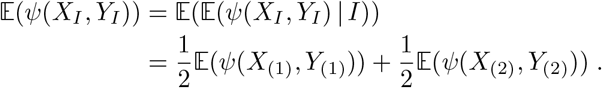

□

#### Proposition 2.

*We have:* 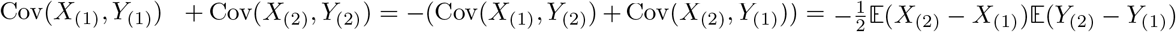.

*Proof*: Consider again the same random index *I*, equal to “(1)” or “(2)” each with probability 1*/*2, independent from (*X*_1_, *Y*_1_, *X*_2_, *Y*_2_). The couples (*X*_*I*_, *Y*_*I*_) and (*X*_*I*_, *Y*_3−*I*_) are both distributed as (*X*_1_, *Y*_1_). Therefore their covariances are null. These covariances can also be computed by conditioning on *I* (see *e*.*g*. formula (1.1) in [37]). For (*X*_*I*_, *Y*_*I*_): Cov(*X*_*I*_, *Y*_*I*_) = 𝔼(Cov(*X*_*I*_, *Y*_*I*_|*I*))+Cov(𝔼(*X*_*I*_|*I*), 𝔼(*Y*_*I*_|*I*)). On the right-hand side, the first term is: 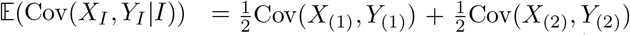. The second term is: 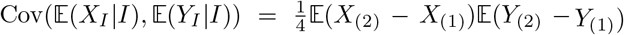. Similarly, we have: Cov(*X*_*I*_, *Y*_3−*I*_) = 𝔼(Cov(*X*_*I*_, *Y*_3−*I*_ |*I*))+Cov(𝔼(*X*_*I*_ |*I*), 𝔼(*Y*_3−*I*_ |*I*)). The first term in the right-hand side is: 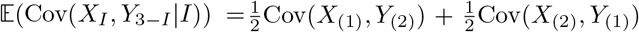. The second term in the right-hand side is: 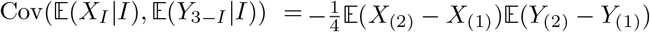. Hence the result. □

#### Proposition 3.

*Assume that the ranking function ϕ is symmetric: ϕ*(*x, y*) = *ϕ*(*y, x*). *Then the couple* (*X*_(1)_, *Y*_(2)_) *has the same distribution as the couple* (*Y*_(1)_, *X*_(2)_).

As a consequence, *X*_(1)_ and *Y*_(1)_ have the same distribution, so do *X*_(2)_ and *Y*_(2)_. Thus: 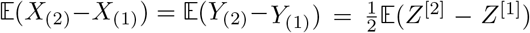. Another consequence is that: Cov(*X*_(1)_, *Y*_(2)_) = Cov(*X*_(2)_, *Y*_(1)_). Thus by Proposition 2: 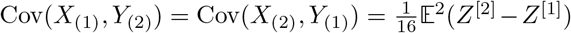.

*Proof*: Since *ϕ* is symmetric, the change of variable (*X*_1_, *Y*_1_, *X*_2_, *Y*_2_) ⟼ (*Y*_1_, *X*_1_, *Y*_2_, *X*_2_) leaves unchanged the couple (*S*_1_, *S*_2_). □

#### Proposition 4.

*Assume that the ranking function ϕ is the sum: ϕ*(*x, y*) = *x* + *y. Then:* 𝔼(*X*_(1)_) = 𝔼(*Y*_(1)_), 𝔼(*X*_(2)_) = 𝔼(*Y*_(2)_), *and* 𝔼(*X*_(1)_) < 𝔼(*X*_(2)_).

*Proof*: The first two equalities come from Proposition 3. By definition, 𝔼(*X*_(1)_ + *Y*_(1)_) < 𝔼(*X*_(2)_ + *Y*_(2)_). Hence the inequality. □

#### Proposition 5.

*Assume that the ranking function ϕ is the sum, and that the common distribution of X*_1_, *Y*_1_, *X*_2_, *Y*_2_ *is symmetric: there exists a such that f* (*x* − *a*) = *f* (*a* − *x*). *Then* (*a* − *X*_(1)_, *a* − *Y*_(1)_) *has the same distribution as* (*X*_(2)_ − *a, Y*_(2)_ − *a*).

As a consequence, Cov(*X*_(1)_, *Y*_(1)_) = Cov(X_(2)_, Y_(2)_).

*Proof*: The change of variable (*X*_1_, *Y*_1_, *X*_2_, *Y*_2_) ⟼ (2*a* − *X*_1_, 2*a*− *Y*_1_, 2*a* − *X*_2_, 2*a* − *Y*_2_) leaves the distribution unchanged. It only swaps the indices (1) and (2) of minimal and maximal sum. □

If we summarize Propositions 1, 2, 3, 4, 5 for the case where the ranking function is the sum, and the distribution is symmetric, one gets:

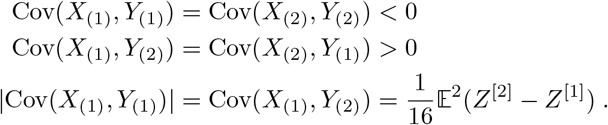

### Two loci, *n* alleles

As in the *n* = 2 case developed above, we are again interested in the statistical properties of the genotypes of maximum fitness. If populations consist of *n* genotypes, the maximal fitness genotypes will have total fitness denoted by random variable *Z*^[*n*]^, the top order statistic of total fitness *Z*. We are more interested, however, in the *concomitants* of *Z*^[*n*]^, namely, random variables *X*_(*n*)_ and *Y*_(*n*)_, defined by the relation *Z*^[*n*]^ = *ϕ*(*X*_(*n*)_, *Y*_(*n*)_). In particular, we are interested in the covariance between the concomitants, cov(*X*_(*n*)_, *Y*_(*n*)_), because changing the sign of this value gives the selective advantage of recombinants (see ev2 [2]).

Before analyzing concomitants of the top order statistic, however, the first step is to derive a general relation between a random variable *Z* and its concomitants *X* and *Y* when these concomitants are defined as *X* + *Y* = *Z*.

#### General relation between Z and its summand concomitants X and Y

Let *n* be an integer larger than 1. For *i* = 1, …, *n*, let (*X*_*i*_, *Y*_*i*_) be i.i.d. couples of random variables. For *i* = 1, …, *n*, let *Z*_*i*_ = *X*_*i*_ + *Y*_*i*_.

Let *U* be a random variable, independent from (*X*_*i*_, *Y*_*i*_), *i* = 1, …, *n*, uniformly distributed over (0, 1). Define the random index *I* in {1, …, *n*} as:

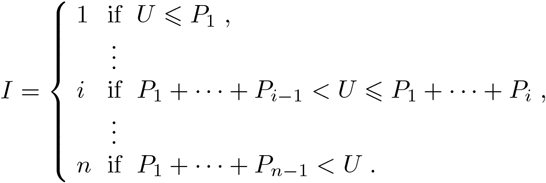

The *P*_*i*_ thus define the discrete fitness distribution governing *Z*. Finally, let (*X, Y*) = (*X*_*I*_, *Y*_*I*_). The goal is to derive statistical properties of concomitants *X* and *Y* of random variable *Z* = *X* + *Y*.

For this, conditioning over two embedded *σ*-algebras, denoted by *ℱ*_2*n*_ and *ℱ*_*n*_, will be used.

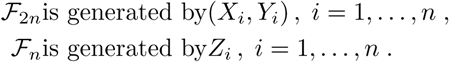

If *A* is any random variable:

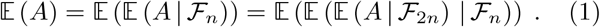

Conditioning functions of (*X, Y*) over ℱ_2*n*_ and ℱ_*n*_ works as follows.

##### Lemma 1.

*Let ϕ be any real valued function of two variables. Provided the following expectations exist, one has:*

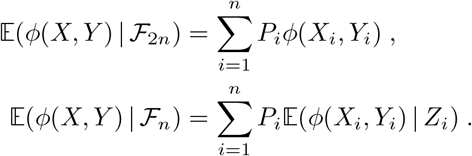

For second order moments, the following well known lemma on conditional covariances will be used.

##### Lemma 2.

*Let* (*A, B*) *be a pair of real-valued random variables on* (Ω, ℱ,ℙ), *and let* ℱ_1_ ⊆ ℱ_2_ *be two σ-fields on* Ω. *Then:*

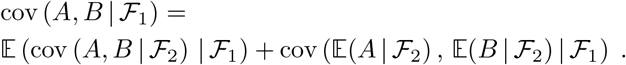

*In particular, when* ℱ_1_ = {Ø, Ω}:

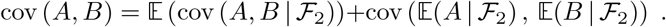

Lemma 3 relates the moments of *X* + *Y* to the *Z*_*i*_’s and *P*_*i*_’s.

##### Lemma 3.

*Denote by* 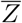 *and V the mean and variance of Z with respect to P:*

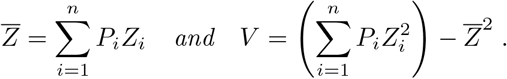

*Then:*

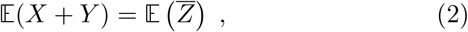

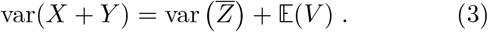

*Proof*: It turns out that 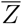 is the conditional expectation of *X* + *Y* with respect to ℱ_*n*_, because:

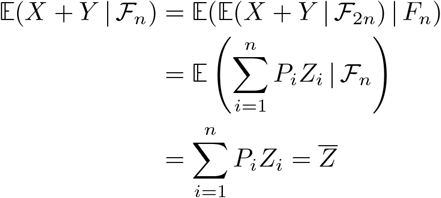

Hence: 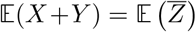. Similarly, *V* is the conditional variance of *X* + *Y*, given ℱ_*n*_. By Lemma 2:

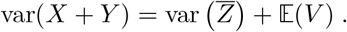

□

From now on, it will be assumed that the common distribution of (*X*_*i*_, *Y*_*i*_), for *i* = 1, …, *n*, is bivariate normal. Lemma 4. *Let* (*X*_1_, *Y*_1_) *be a couple of random variables, having bivariate normal distribution* 𝒩_2_(*μ, K*), *with expectation μ* = (*μ*_*x*_, *μ*_*y*_), *covariance matrix:*

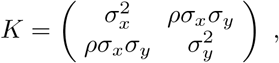

*where σ*_*x*_ > 0, *σ*_*y*_ > 0, |*ρ*| < 1.

*Denote:*

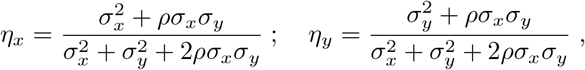

*and also:*

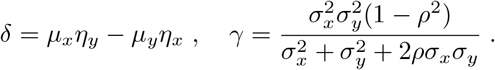

*Let Z*_1_ = *X*_1_ + *Y*_1_. *The conditional distribution of* (*X*_1_, *Y*_1_) *given Z*_1_ = *z is bivariate normal, with expectation:*

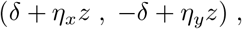

*covariance matrix:*

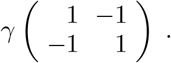

*Proof*: The vector (*X*_1_, *Y*_1_, *Z*_1_) has normal distribution with expectation (*μ*_*x*_, *μ*_*y*_, *μ*_*x*_+*μ*_*y*_), and covariance matrix:

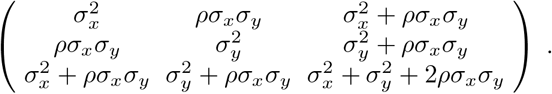

The conditional distribution of (*X*_1_, *Y*_1_) given *Z*_1_ = *z* is again normal. The conditional expectation of *X*_1_ is:

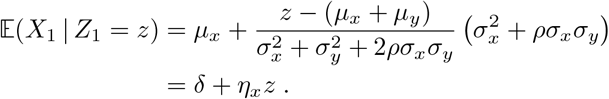

The conditional expectation of *Y*_1_ is symmetric:

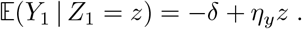

The covariance matrix does not depend on *z*:

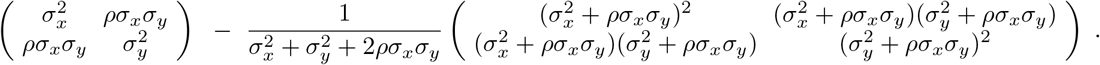

After simplification one gets:

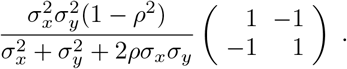

□

Theorem 1 below gives the first and second order moments of the random couple (*X, Y*), when the common distribution of the (*X*_*i*_, *Y*_*i*_) is that of Lemma 4.

##### Theorem 1.

*Assume that for i* = 1, …, *n, the distribution of* (*X*_*i*_, *Y*_*i*_) *is bivariate normal* 𝒩_2_(*μ, K*). *With the notations of Lemma 4:*

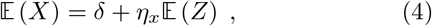

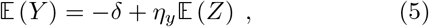

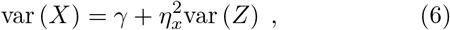

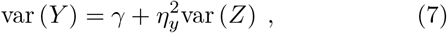

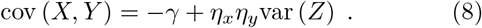

Observe that, since *η*_*x*_ + *η*_*y*_ = 1, the first two equations add to identity, and so do the last three, the last one being doubled.

*Proof*: By Lemma 1,

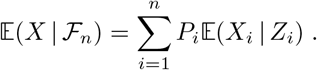

By Lemma 4,

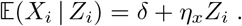

Hence:

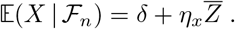

Similarly:

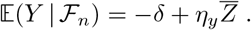

Let us now compute var(*X*). By Lemma 2:

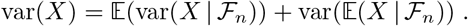

By Lemma 1,

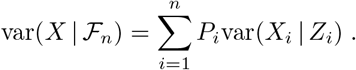

But by Lemma 4, var(*X*_*i*_|*Z*_*i*_) is the constant *γ*, independently on *Z*_*i*_. Thus:

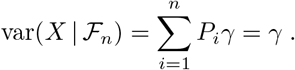

Now by Lemma 1:

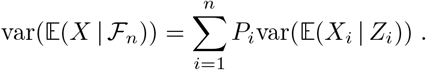

By Lemma 4, 𝔼(*X*_*i*_ | *Z*_*i*_) = *δ* + *η*_*x*_*Z*_*i*_, hence:

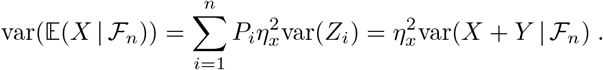

Joining both results through Lemma 2:

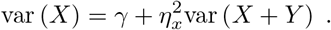

Similarly:

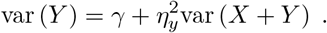

Let us now turn to cov(*X, Y*): By Lemma 2:

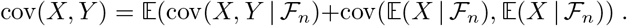

By Lemma 1,

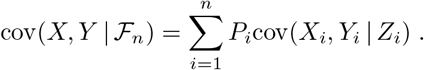

But by Lemma 4, cov(*X*_*i*_, *Y*_*i*_ *Z*_*i*_) is the constant *γ*, independently on *Z*_*i*_. Thus:

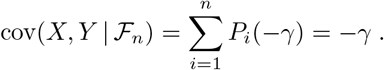

Now by Lemma 1:

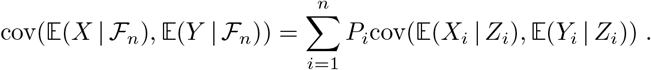

By Lemma 4, 𝔼(*X*_*i*_ | *Z*_*i*_) = *δ* + *η*_*x*_*Z*_*i*_, and 𝔼(*Y*_*i*_ | *Z*_*i*_) = −*δ* + *η*_*y*_*Z*_*i*._ Hence:

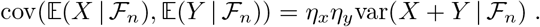

Joining both results through Lemma 2:

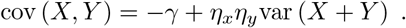

□

We define new random variable

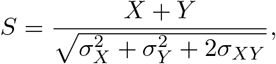

which is 𝒩(0, 1). The *k*^*th*^ order statistic of *S* is denoted *S*^[*k*]^ and is related to its concomitant summands as:

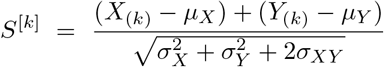

[38]. Rearranging gives:

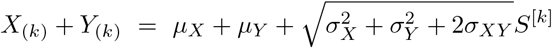

from which we have:

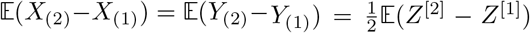

Plugging this expression into Eqs (6), (7) and (8) leads to the following corollary.

##### Corollary 1.

*Define random variable S* ∼ 𝒩(0, 1) *whose k*^*th*^ *order statistic from a sample of size n is denoted S*^[*k*]^. *If we blindly (and wrongly) assume that order statistic distributions are normal, the first- and second-order moments of the concomitants are nevertheless exact:*

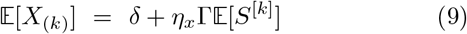

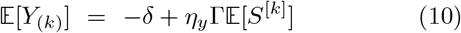

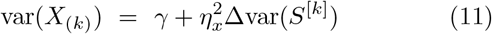

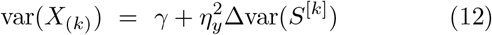

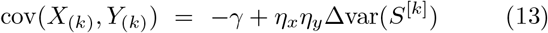

*where* 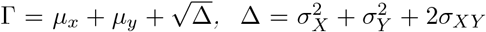, *and X*_(*k*)_ + *Y*_(*k*)_ = *Z*^[*k*]^, *the k*^*th*^ *order statistic in total fitness*.

Comparison with simulations show the foregoing expressions to be extremely accurate. And comparison with previous studies that take a more circuitous route in different contexts [39–41] show these expressions to be exact. While our analysis is more compact than those previous studies, we suspect our approach would not be exact for higher moments.

In general, we are most interested in the top order statistic, *k* = *n*, because natural selection will tend to “select” the fittest genotype. In an infinite population, selection is completely deterministic and the fittest will always be fixed. We note that our findings are only weakly dependent on this assumption of deterministic selection, because the variance of top order statistics are often quite similar; hence, our findings remain relatively unchanged if suboptimal genotypes *k* = *n*−1 or *k* = *n*−2 are selected due to finite-population effects (drift). We show in [2] that the mean selective advantage of recombinants will be:

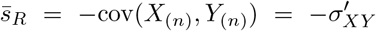

where the final step is simply a change of notation. We define 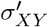 to be the covariance of the concomitants of the top order statistic, i.e., the covariance between *X* and *Y* after natural selection has acted locally in each of the subpopulations; *σ*_*XY*_ retains its meaning as general covariance (not covariance of concomitants) between *X* and *Y* in the initial population. More simply, *σ*_*XY*_ is preselection covariance and 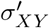 is post-selection covariance.

We further define the variance of the top order statistic of a standard normal random variable: 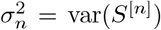, which has the property 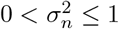 [38, 42]. Full general expressions are given in the SM. Two simplified cases are illuminating and are discussed here:

The first illuminating case is when *X* and *Y* are independent. In this case, Eq (13) becomes:

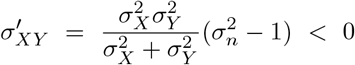

because two or more genotypes (*n* ≥ 2) are required for recombination to make a difference, and 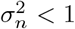 for *n* ≥ 2. In words, after natural selection has run its course in local subpopulations, recombination across those local subpopulations will be advantageous.

The second illuminating case is when *σ*_*X*_ = *σ*_*Y*_ = *σ*. In this case, we have the following equivalent expressions:

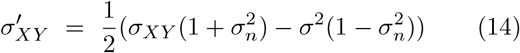

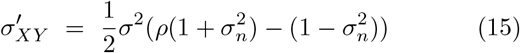

where *ρ* = *σ*_*XY*_ (*σ*_*X*_ *σ*_*Y*_)^−1^, the pre-selection correlation coefficient.

The first thing to notice is that the effect of natural selection is always to reduce covariance by an amount whose lower bound depends only on the number of competing genotypes:

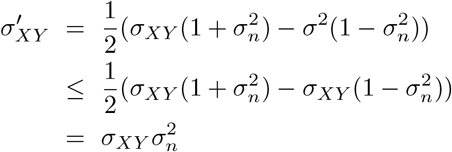

In the above expressions, it is apparent that post-selection covariance can in theory be positive if pre-selection covariance is strongly positive. (In other words, post-selection recombinant advantage can in theory be negative if pre-selection recombinant advantage is strongly negative.) The condition for covariance to be negative after a single bout of selection is best expressed as a condition on pre-selection correlation; post-selection covariance will be negative when:

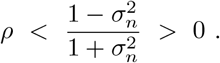

In theory, at least, one bout of selection may not result in negative covariance. Several bouts of selection, however, are guaranteed to result in negative covariance. The equilibrium covariance achieved after many bouts of selection can be determined by setting 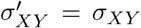 in Eq (14), giving:

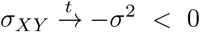

For the general case where the variances are not equal,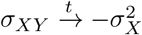 and 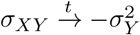 are both stable equilibria.

## V. HETEROSIS

Our findings can be viewed as providing a theoretical basis for a kind of haploid heterosis. We will now show that our findings also provide a theoretical basis for classical diploid heterosis as well. After the rediscovery of Mendel’s work, two competing mechanistic explanations for heterosis emerged:

The first explanation – the *dominance hypothesis* – relied on two observations: 1) inbreeding tends to produce homozygotes, and 2) deleterious alleles tend to be recessive. If a locus is homozygous dominant (wildtype) in one population and homozygous recessive (deleterious) in the other, an across-population recombination event has probability 1/4 of producing a deleterious offspring, whereas it would have probability 3/4 in the absence of dominance.

The second explanation – the *overdominance hypothesis* relied on empirical observations of a curious phenomenon (overdominance) [14, 16, 44, 48, 49], where heterozygotes at a given locus are fitter than either homozygote. While overdominance has been observed, and there are famous examples, the genetic/mechanistic basis of overdominance is varied and nebulous.

Increasingly detailed studies of heterosis reveal that observations of overdominance are not really overdominance at all but are in fact an artefact of linkage that can give the *appearance* of overdominance [50–52]. In these cases, what appears to be a single locus is in fact two or more loci in linkage with each other. If the alleles at the linked loci are selectively mismatched giving rise to negative associations in genic fitnesses across populations, out-crossing between populations can give the appearance of overdominance as fitter dominants mask less-fit recessives – a phenomenon that has been dubbed *pseudo-overdominance* [45, 50–53]. (What we are here calling “selective mismatch” has elsewhere been called “linkage repulsion” [52–54], “linkage bias” [51], or “linkage disequilibrium” (LD) [47].) If the out-crossed parents come from different inbred populations – a common practice in agriculture – they will have high homozygosity and the heterosis effect in the offspring will be accentuated; a schematic of this scenario is presented in Fig 6.

The weak link in the pseudo-overdominance theory is the requirement that linkage repulsion (selective mismatch) somehow develop within blocks of linked loci – dubbed pseudo-overdominance blocks (or PODs) [51, 52]. Some authors have made verbal arguments invoking a combination of mutation, weak mutational effect and small effective population size to meet this requirement [43]. In a very recent paper, Waller [52] makes an elegant argument showing how slightly-deleterious mutations can mask strongly-deleterious mutations, thereby maintaining inbreeding depression over long periods of time – an observation that confounded Darwin. The theory we have developed in the present study speaks directly to the requirement of linkage repulsion and provides a general theoretical foundation for the pseudo-overdominance theory of heterosis. Mutation, weak mutational effects and small population sizes are not required. The required linkage repulsion is produced across populations of any size simply by natural selection acting on heritable variation. Conceptually, heterosis due to pseudo-overdominance can ultimately be a product of the counter-intuitive phenomenon outlined in Figs 2 and 3 of our companion paper [1].

**Figure 3.**
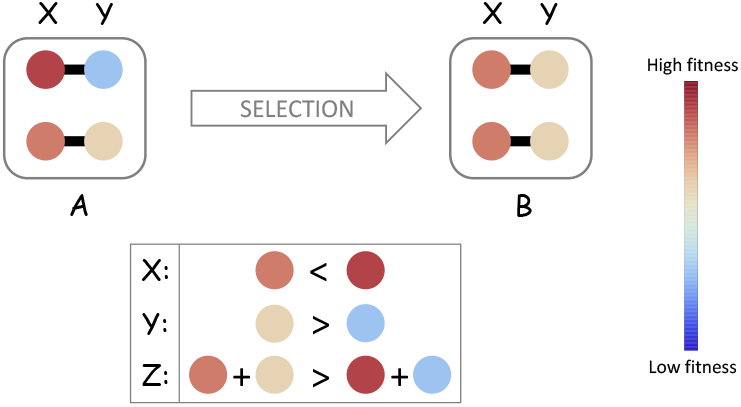
Two loci, two alleles. Here, a large (infinite) population consists of individuals whose genome has only two loci *x* and *y*, each of which carries one of two alleles: genotype 1 carries allele *X*_1_ at the *x* locus and *Y*_1_ at the *y* locus, and genotype 2 carries allele *X*_2_ at the *x* locus and *Y*_2_ at the *y* locus. An individual’s fitness is simply the sum of its genic fitnesses, *Z* = *X* + *Y*, so that the fitnesses of genotypes 1 and 2 are *Z*_1_ = *X*_1_ + *Y*_1_ and *Z*_2_ = *X*_2_ + *Y*_2_, respectively. The fitter of these two genotypes has total fitness denoted *Z*^[2]^ (i.e., *Z*^[2]^ = Max{*Z*_1_, *Z*_2_}) and genic fitnesses *X*_(2)_ and *Y*_(2)_ (i.e., *Z*^[2]^ = *X*_(2)_ + *Y*_(2)_). Similarly, the less-fit of these two genotypes has total fitness *Z*^[1]^ = *X*_(1)_ +*Y*_(1)_. We note: *Z*^[2]^ > *Z*^[1]^ by definition, but this does *not* guarantee that *X*_(2)_ > *X*_(1)_ or that *Y*_(2)_ > *Y*_(1)_, as illustrated in the lower box. The population labeled *A* consists of two distinct genotypes but selection acts to remove the inferior genotype leaving a homogeneous population in which individuals are all genetically identical (with fitness *Z*^[2]^) as illustrated in the population labeled *B*.

**Figure 4.**
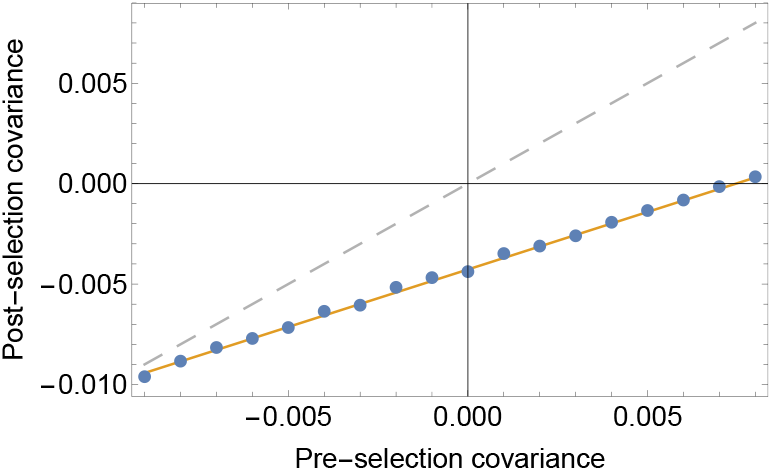
Covariance before and after selection. Blue dots plot covariance across simulations of 5000 subpopulations, each containing *n* = 20 distinct genotypes and a bivariate normal distribution with means equal to −0.1, variances equal to 0.01, and covariance indicated on the horizontal axis. Orange line plots theoretical prediction given by Eq (14). Gray dashed line plots *y* = *x* as a visual guide. Post-selection covariance is suppressed more when pre-selection covariance is strongly positive.

**Figure 5.**
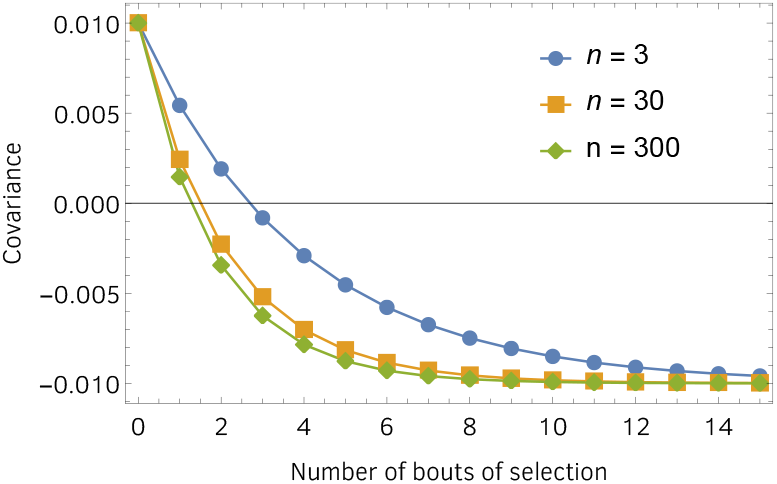
Covariance dynamics over several bouts of selection. Initially, we assign covariance its maximal possible value of *σ*_*X*_ *σ*_*Y*_ = 0.01 in order to illustrate the fact that, at least in theory (under an extreme condition), it may take more than one bout of selection for covariance to become negative. Covariance does become negative rather quickly, however, and converges to the predicted value of −*σ*_*X*_ *σ*_*Y*_ = −0.01, which is the minimal value for covariance.

**Figure 6.**
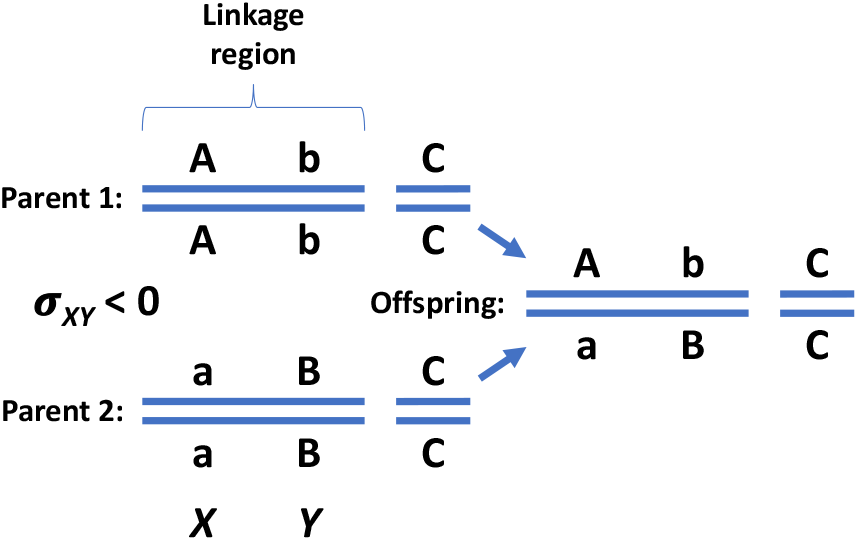
A novel theoretical basis for pseudo-overdominance and heterosis. Upper case (lower case) letters denote higher- (lower-) fitness alleles. Generally speaking, lower-fitness alleles tend to be recessive. Heterozygotes will therefore tend to express the higher-fitness of the two alleles at a locus. Here, parents 1 and 2 come from two different inbred populations. Inbreeding populations tend toward homozygosity and our findings show that natural selection will tend to fix alleles that are poorly matched across loci, i.e., that have negative covariance in genic fitnesses. In the simple example illustrated here, the parents expressing phenotypes (A,b,C) and (a,B,C) exhibit negative covariance in genic fitness across loci. Both parents are less fit that the offspring which expresses phenotype (A,B,C), thereby generating heterosis through an appearance of overdominance (called *pseudo-overdominance*) [43–47]. Our theory shows how the required negative covariance in genic fitnesses across linked loci is an unavoidable consequence of natural selection, and thus provides a novel theoretical basis for heterosis.

## VI. CONCLUDING REMARKS

To summarize what has been modeled in this paper, we revisit our definition of “products of selection”. These are genotypes that are locally prevalent, due to natural selection. Products of selection can include locally-prevalent genotypes in populations, subpopulations, demes, niches, or competing clones. A spatially-structured population, for example, can have many spatially separated subpopulations. After selection has been operating in these sub-populations for some time, if an individual from one sub-population recombines with an individal from another subpopulation, our findings show that the offspring will be fitter, on average, than both parents.

In simulations (SM), we placed such recombinant offspring in head-to-head competition with non-recombinant offspring, with no further recombination occurring during the competition. We found that the recombinant offspring displaced the non-recombinant off-spring > 95% of the time under a wide range of conditions.

The mathematical analyses in this study is a bit more restrictive than our analyses in companion paper [2], most of which has zero dependence on the initial parent distribution of genic fitnesses. Here, our mathematical analyses eventually require: 1) an assumption that the initial distribution is symmetric, for the 2-locus, 2-genotype case, and 2) an assumption that the initial distribution is normal, for the 2-locus, *n*-genotype case. Our finding that the lower central moments of concomitants *X*_(*k*)_ and *Y*_(*k*)_ are exact despite a bold assumption of normality is at least suggestive that our findings might be robust more generally, i.e., to non-normal parent distributions. Furthermore, our qualitative results are corroborated by simulations with a wide variety of divergent parent distributions (SM).

Finally, our findings correct the straw-man argument, outlined in the first paragraph of this paper, commonly used to demonstrate why sex and recombination are enigmatic. The premise of this argument is that natural selection will tend to amplify genotypes that carry “good” (selectively well-matched) combinations of genes. We find that, when natural selection is operating in isolation, the opposite is true quite generally. We find that natural selection has an encompassing tendency to amplify genotypes carrying “bad” (selectively mis-matched) combinations of genes. Recombination on average breaks up bad combinations and assembles good combinations, and its evolution is thus promoted.

## ACKNOWLEDGEMENTS

Much of this work was performed during a CNRS-funded visit (P.G.) to the Laboratoire Jean Kuntzmann, University of Grenoble Alpes, France, and during a visit to Bielefeld University (P.G.) funded by Deutsche Forschungsgemeinschaft (German Research Foundation, DFG) via Priority Programme SPP 1590 Probabilistic Structures in Evolution, grants BA 2469/5-2 and WA 967/4-2. P.G. and A.C. received financial support from the USA/Brazil Fulbright scholar program. P.G. and P.S. received financial support from National Aeronautics and Space Administration grant NNA15BB04A. P.G. received further support from the National Institute Of General Medical Sciences of the National Institutes of Health under Award Number R35GM137919 (awarded to Gideon Bradburd). The authors thank S. Otto and N. Barton for their thoughts on early stages of this work. Special thanks go to E. Baake for her thoughts on later stages of this work and help with key mathematical aspects. The authors thank D. Chencha, J. Streelman, R. Rosenzweig and the Biology Department at Georgia Institute of Technology for critical infrastructure and computational support.

